# Loss of Neutralizing Antibody Response to mRNA Vaccination against SARS-CoV-2 Variants: Differing Kinetics and Strong Boosting by Breakthrough Infection

**DOI:** 10.1101/2021.12.06.471455

**Authors:** John P. Evans, Cong Zeng, Claire Carlin, Gerard Lozanski, Linda J. Saif, Eugene M. Oltz, Richard J. Gumina, Shan-Lu Liu

## Abstract

The waning efficacy of SARS-CoV-2 vaccines combined with the continued emergence of variants resistant to vaccine-induced immunity has reignited debate over the need for booster vaccines. To address this, we examined the neutralizing antibody (nAb) response against four major SARS-CoV-2 variants—D614G, Alpha (B.1.1.7), Beta (B.1.351), and Delta (B.1.617.2)—in health care workers (HCWs) at pre-vaccination, post-first and post-second mRNA vaccine dose, and six months post-second mRNA vaccine dose. Neutralizing antibody titers against all variants, especially the Delta variant, declined dramatically from four weeks to six months post-second mRNA vaccine dose. Notably, SARS-CoV-2 infection enhanced vaccine durability, and mRNA-1273 vaccinated HCWs also exhibited ~2-fold higher nAb titers than BNT162b2 vaccinated HCWs. Together these results demonstrate possible waning of protection from infection against SARS-CoV-2 Delta variant based on decreased nAb titers, dependent on COVID-19 status and the mRNA vaccine received.

## Introduction

Since its emergence in late 2019, the COVID-19 pandemic has led to over 252 million confirmed cases and over 5 million deaths as of November 14, 2021 (1). In response, several vaccines have been developed against SARS-CoV-2, the causative agent of COVID-19, including two novel mRNA vaccines, Moderna mRNA-1273 and Pfizer/BioNTech BNT162b2. These highly effective vaccines have helped to stem COVID-19 hospitalizations and deaths. However, the rapid evolution of SARS-CoV-2, combined with waning vaccine efficacy, remain a threat to public health.

Following its introduction into the human population, several SARS-CoV-2 variants of concern (VOCs) have emerged. Very soon after zoonotic transmission, SARS-CoV-2 acquired a predominant D614G mutation in its spike (S) protein. This mutation leads to enhanced transmissibility, likely due to increased stability of the S protein, increased viral titers in the nasopharynx, and increased infectivity (2). As a result, nearly all currently circulating SARS-CoV-2 strains bear the D614G mutation (3). However, as greater proportions of the world population acquired immunity against SARS-CoV-2, through infection or vaccination, new VOCs emerged that had reduced susceptibility to antibody-mediated immune responses and continued to become more transmissible (4, 5). One VOC, Alpha (B.1.1.7), is characterized by N-terminal domain (NTD) deletions and a key N501Y mutation in its receptor-binding domain (RBD). Alpha exhibited enhanced transmissibility and rapidly spread from Europe to other parts of the world (6). Another VOC to emerge at about the same time was Beta (B.1.351), which is characterized by other NTD mutations and deletions, as well as key RBD mutations, including K417N, E484K, and N501Y. While the Beta variant did not disseminate as widely as Alpha, it harbored strong resistance to vaccine-induced immunity (7). Finally, Delta (B.1.617.2) is responsible for the most recent wave of the COVID-19 pandemic and is characterized by new NTD alterations, together with key RBD mutations (L452R and T478K). Delta has led to an alarming number of vaccine breakthrough infections worldwide and has prompted debate about the need for vaccine booster doses.

The extent to which the rise in breakthrough infections is caused by increased resistance to vaccine-induced immunity in these variants and/or to waning durability of immunity and efficacy of vaccines in preventing infection remains unclear. Reports from India, where the population was still pursuing mass vaccination efforts, show minor differences in breakthrough infection rates between Alpha and Delta. Specifically, BNT162b2 efficacy against symptomatic infection was reported to drop from 93.4% against Alpha to 87.9% against Delta (8). However, reports from the U.S. indicate that vaccine efficacy of BNT162b2 against Delta infection declined from 93% one month after vaccination to 53% at four months (9), consistent with an overall waning of vaccine efficacy over time (10). A critical goal of this study is to better understand how the durability of vaccine efficacy contributes to rates of breakthrough infections, especially in the context of evolving SARS-CoV-2 variants. Such insights will improve strategies for allocation of booster doses, recommendations for immunocompromised patients, and could guide any reformulation of future SARS-CoV-2 booster doses.

To address these issues, we examined neutralizing antibody (nAb) levels in 48 vaccinated health care workers (HCWs) against the major SARS-CoV-2 variants using serum collected prevaccination, one month after the first dose of BNT162b2 or mRNA-1273, and one and six months after the second dose of vaccine. Indeed, prior studies have shown that neutralizing antibody levels are a major correlate for protection from SARS-CoV-2 infection (11).

## Results

We produced lentiviral pseudotypes expressing a *Gaussia* luciferase reporter gene and bearing SARS-CoV-2 spike derived from D614G, Alpha, Beta, or Delta (**Fig. 1A**). Pseudotyped virus infectivity was then determined by infection of HEK293T-ACE2 cells. *Gaussia* luciferase secreted into the media of infected cells was assayed to determine the infectivity of produced lentiviral pseudotypes. We did not find significant differences in pseudotyped lentivirus infectivity for the four variants (all containing D614G) tested (**Fig. 1B**), despite some reports of drastically increased transmission and spread for some VOCs, especially the Delta variant (12).

**Figure 1:**
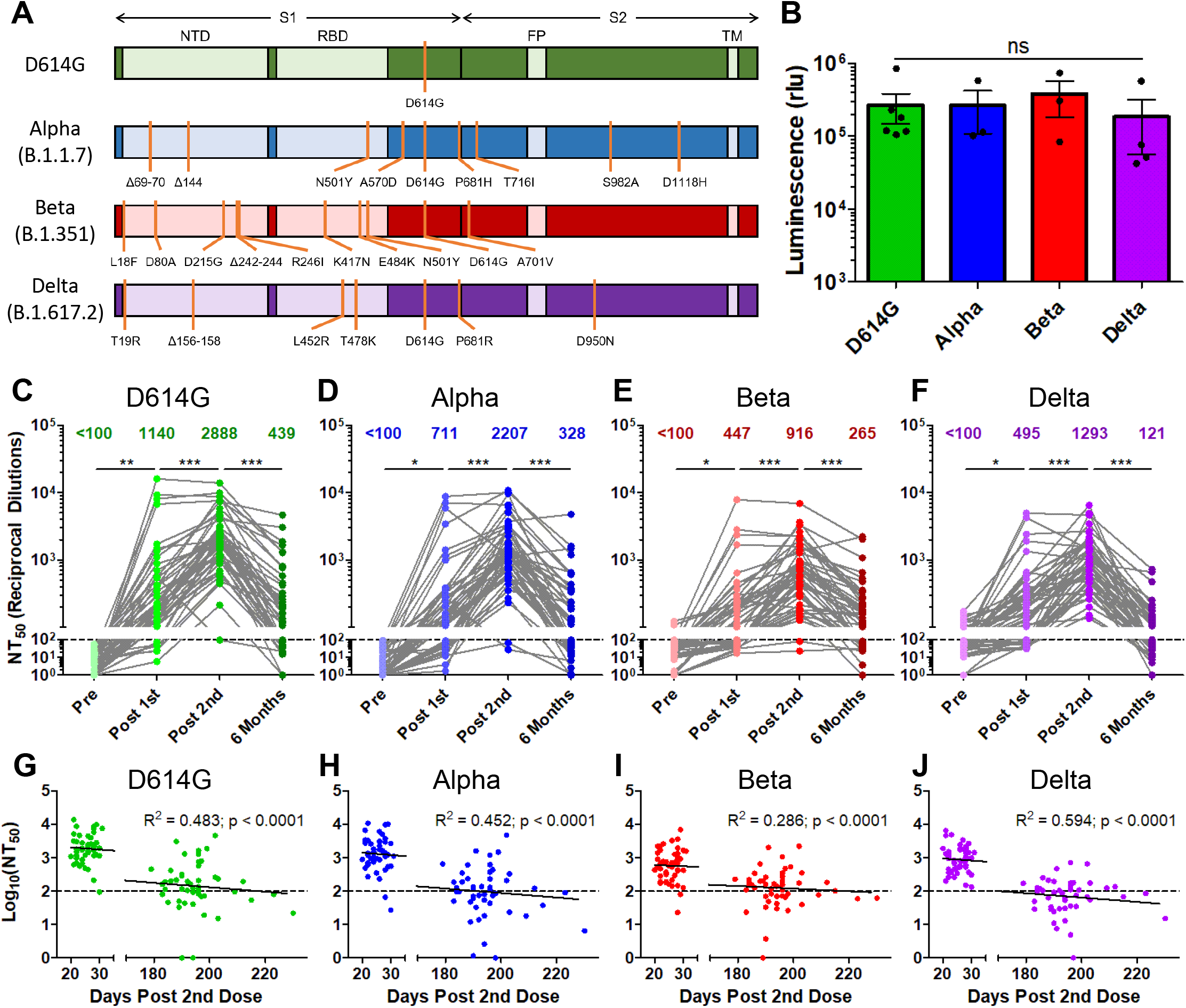
The durability of vaccine-induced immunity wanes over time, with a virtual loss at six months for the Delta variant. *Gaussia* luciferase reporter gene containing lentivirus pseudotypes were produced bearing the spike (S) protein from SARS-CoV-2 variants. (**A**) Schematic representations of the SARS-CoV-2 variant spikes tested are shown which contain the indicated mutations. These include D614G, Alpha (B.1.1.7), Beta (B.1.351), and Delta (B.1.617.2). The schematics highlight the location of the S1 and S2 subunits as well as the N-terminal domain (NTD), receptor binding domain (RBD), fusion peptide (FP), and transmembrane region (TM). (**B**) Lentivirus pseudotypes were used to infect HEK293T-ACE2 cells and 48hrs after infection media was harvested from infected cells and assayed for *Gaussia* luciferase activity to determine the relative infectivity of variant pseudotyped virus. (**C-F**) Lentivirus pseudotyped with SARS-CoV-2 S from D614G (**C**), Alpha (**D**), Beta (**E**), and Delta (**F**) were incubated for 1 hr to neutralize with serial dilutions (1:80, 1:320, 1:1280, 1:5120, 1:20480, and no serum) of health care worker (HCW) serum collected pre vaccination, post vaccination with a first dose of Pfizer/BioNTech BNT162b2 or Moderna mRNA-1273, post vaccination with a second dose of mRNA vaccine, and six months post vaccination with a second dose of mRNA vaccine. Neutralized virus was then used to infect HEK293T-ACE2 cells and *Gaussia* luciferase activity was assayed 48 hrs and 72 hrs after infection. Neutralization titers 50% (NT_50_) were determined by least-squares fit non-linear regression. Mean NT_50_ are shown at the top of the plots, and NT_50_ values below 100 were considered background. (**G-J**) Log_10_ transformed (to better approximate linearity) NT_50_ values against D614G (**G**), Alpha (**H**), Beta (**I**), and Delta (**J**) variants were plotted against days post-second vaccine dose of sample collection. The equation of the fitted curve, the goodness of fit (R^2^), and p-value for the curve are displayed on each plot. The dotted lines correspond to the background level (NT_50_ < 100). Statistical significance was determined by one-way ANOVA with Bonferroni’s correction (**B**), one-way repeated measures ANOVA with Bonferroni’s correction (**C-F**), or by least-squares fit linear regression (**G-J**). In call cases, *p < 0.05; **p < 0.01; ***p < 0.001; ns: not significant.

We used our previously reported (13, 14) highly-sensitive SARS-CoV-2 pseudotyped lentivirus-based virus neutralization assay to assess nAb titers in HCW samples collected under approved IRB protocols (2020H0228 and 2020H0527). The 48 HCW samples included 22 mRNA-1273 and 26 BNT162b2 vaccinated individuals, with a median age of 37 years (IQR = 31.75-43.25). Samples were collected from HCWs with median time points of 222 days (IQR = 215-225.75) pre-first vaccine dose (Pre), 21 days (IQR = 19.25-23) post-first vaccine dose (Post 1st), 26 days (IQR = 22.5-28) post-second vaccine dose (Post 2nd), and 194 days (IQR = 190=197.75) post-second vaccine dose (Six Months). According to the titer of pseudotyped viruses, we adjusted the volumes of each so that equivalent infectious viruses were used in neutralization assays. HCW serum samples underwent 4-fold serial dilutions followed by the addition of pseudotyped virus for one hr neutralization, with final dilutions of 1:80, 1:320, 1:1280, 1:5120, 1:20480, and no serum control. HEK293T-ACE2 cells were then infected with neutralized virus and *Gaussia* luciferase activity was assayed 48 hrs and 72 hrs after infection. Neutralizing titer 50% (NT_50_) values were determined by least-squares fit, non-linear regression in GraphPad Prism 5.

We compared the strength of the nAb titers over time against all four variants tested. Following the first dose of mRNA vaccine, a strong nAb response was induced among HCWs compared to pre-vaccination across all variants (p < 0.001), which efficiently blocked virus entry; this was despite the huge variation in nAb titers of these individuals including against D614G (mean = 1140, 95% CI = 317-1963, range = 100-15954) (**Fig. 1C**). However, across all variants, between 14.6% (7/48) and 45.8% (22/48) of HCWs exhibited NT_50_ values below detection limit (NT_50_ < 100) following the first dose of vaccine (**Fig. 1C-F**). These initial nAb titers fell to 0.0% (0/48) to 4.2% (2/48) for all variants following a second vaccine dose, with a 2-3-fold increase in mean nAb titers compared to the first dose (p < 0.001) (**Fig. 1C-F**). Notably, four HCWs with higher nAb titers after the first vaccine dose did not show an increase, but a plateau or slight decline in nAb titers following the second dose (**Fig. 1C-F**). These four individuals included one that was anti-SARS-CoV-2-N positive at pre-vaccination, and three that were anti-N positive post-first vaccine dose—indicating infection either prior to or shortly after their first vaccine dose. We found that, following two vaccine doses, the Alpha, Beta, and Delta VOCs exhibited a 1.3- (p < 0.001), 3.2- (p < 0.001), and 2.2-fold (p < 0.001) lower NT_50_ values compared to D614G, respectively (**Fig. 1C-F**). Critically, six months post-vaccination, there was a 3.5-10.7-fold reduction in nAb levels against all variants examined, with 37.5% (18/48) to 56.3% (27/48) of HCWs exhibiting NT_50_ levels below the limit of detection (**Fig. 1C-F**). The mean NT_50_ values for Alpha, Beta, and Delta variants at six months were 1.3-, 1.7-, and 3.6-fold lower than that of D614G, respectively, although the differences in these low nAb titer groups were not statistically significant (**Fig. 1C-F**).

We also examined the correlation between time post-second dose and log_10_ transformed NT_50_ values. We found a statistically significant association between these values for all four variants (**Fig. 1G-J**). This corresponded to an approximately 10-fold decline in NT_50_ for D614G, Alpha, and Delta (R^2^ = 0.0452-0.594, p < 0.0001) every ~22 weeks compared with Beta (R^2^ = 0.286, p < 0.001) every ~37 weeks (**Fig. 1G-J**).

Prior COVID-19 status is a critical parameter for the nAb response to vaccination (15). Of the 48 HCWs examined, one was anti-SARS-CoV-2 N positive by ELISA pre-vaccination, four were anti-N positive at their post-first vaccine dose sample, three at their post-second vaccine dose sample, and four at their six-month vaccine sample—indicating that these 12 subjects were infected by SARS-CoV-2 at different phases of vaccination (**Fig. 2A**). At the time of pre-vaccination sample collection D614G was the major circulating SARS-CoV-2 variant, while at the time off post-first dose and post-second dose D614G and Alpha were circulating, and at the six month time point Delta was the dominant strain. Notably, not all patients remained anti-N positive, but were still considered to have been infected for the purpose of analysis. Following the first vaccine dose, anti-N positive HCWs exhibited 11.7-fold higher mean NT_50_ (p < 0.001) against all four viruses compared to the anti-N negative HCWs (**Fig. 2B**). This difference diminished to 2.3-fold following a second vaccine dose (p < 0.001) (**Fig. 2B**). However, at six months post-vaccination, anti-N positive HCWs exhibited 6.1-fold higher NT_50_ values than anti-N negative HCWs for all variants (p = 0.042) (**Fig. 2B**). Interestingly, we found that the differences in NT_50_ between anti-N positive and negative HCWs were greater and more statistically significant for D614G and Alpha compared with the Beta and Delta variants, likely due to the strong neutralization resistance of the latter VOCs (**Fig. 2C**). Notably, for anti-N negative HCWs, between 41.7% (15/36) and 66.7% (24/36) of subjects exhibited NT_50_ against all four variants that were below detection limit at six months, in sharp contrast to anti-N positive individuals, who were between 8.3% (1/12) and 25.0% (3/12) (**Fig. 2C**).

**Figure 2:**
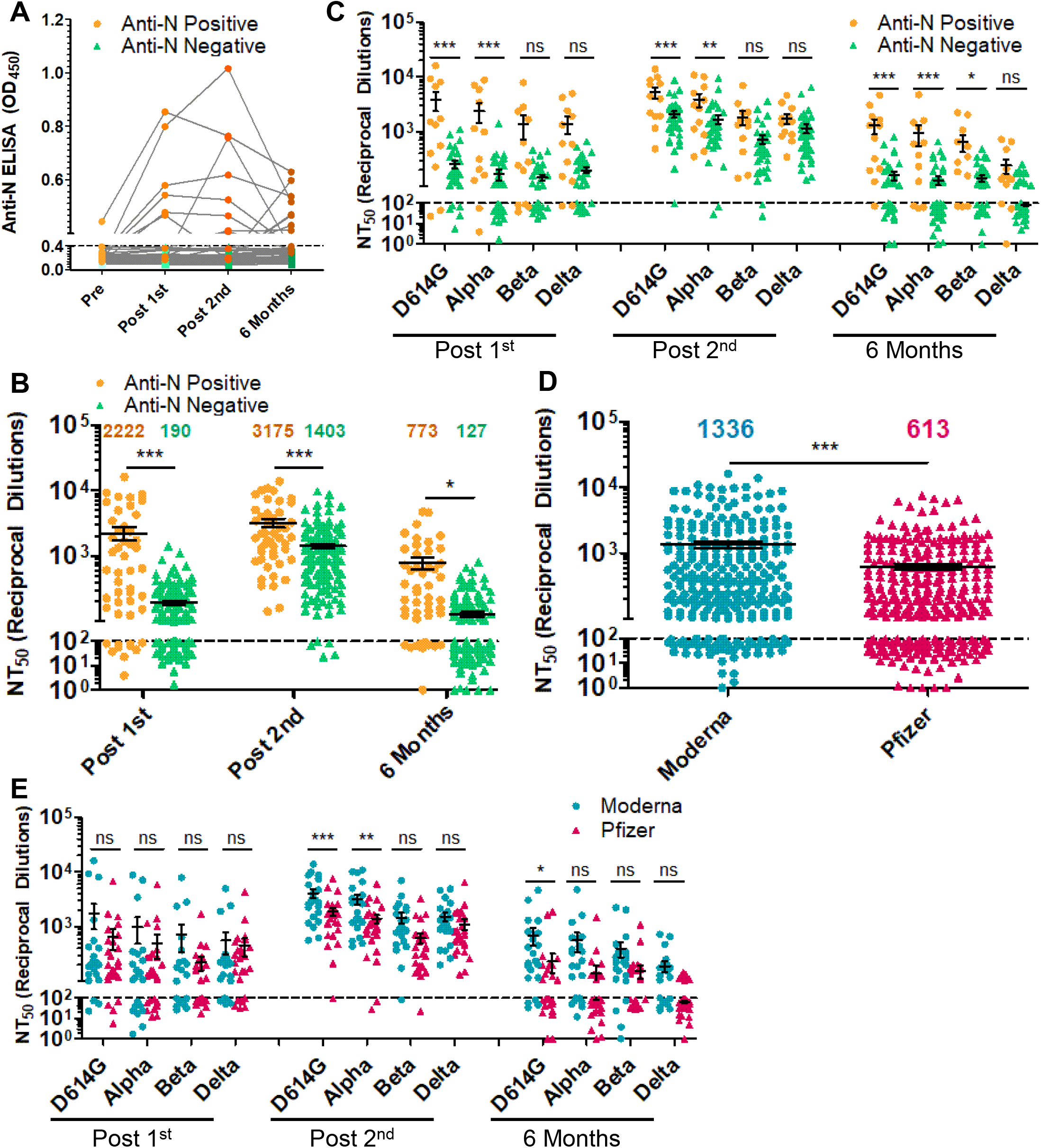
The durability of the nAb response is dependent on prior COVID-19 status, mRNA vaccine type but not age. (**A**) Anti-N ELISA results are presented for HCWs who became anti-N positive (OD_450_ > 0.4 at any time point; n = 12) and HCWs who never became anti-N positive (OD_450_ < 0.4 for all time points; n = 36). (**B, C**) HCWs were divided by prior COVID-19 status as determined by anti-SARS-CoV-2 N ELISA. HCWs with anti-N above the cut-off value of 0.4 for any time point (n = 12) were considered as COVID-19 positive during the study period. NT_50_ values against all four variants combined (**B**) or separated (**C**) for anti-N positive HCWs are compared to anti-N negative HCWs for samples collected post first mRNA vaccine dose, post second mRNA vaccine dose, and six months post second mRNA vaccine dose, respectively. (**D, E**) HCWs were divided by types of mRNA vaccine received, either Moderna mRNA-1273 (n = 22) or Pfizer/BioNTech BNT162b2 (n = 26), and all variants at post-first vaccine dose, post-second vaccine dose, and six months post-second vaccine doses were plotted together (**D**) or grouped by variant and time point (**E**). Mean NT_50_ values are indicated at the top of plots (**B, D**) Statistical significance was determined by one-way ANOVA with Bonferroni’s correction (**B**) two-way, repeated measures ANOVA with Bonferroni’s correction (**C, E**) or unpaired two-tailed t-test with Welch’s correction (**D**). In call cases, *p < 0.05; **p < 0.01; ***p < 0.001; ns: not significant.

We further examined the difference in nAb durability between Moderna mRNA-1273 and Pfizer/BioNTech BNT162b2 vaccinated HCWs. Across all variants over the full-time course, we observed that mRNA-1273 elicited an overall 2.2-fold higher nAb response than the BNT162b2 (p < 0.001) (**Fig. 2D**). In particular, following two vaccine doses, mRNA-1273 vaccinated HCWs exhibited 2.1-, 2.3-, 2.4-, and 1.3-fold higher nAb response compared to BNT162b2-vaccinated HCWs for D614G, Alpha, Beta, and Delta variants, respectively (**Fig. 2E**). The slightly higher NT_50_ values of mRNA-1273 vaccinated HCWs persisted through their six-month collection (**Fig. 2E**), with 18.2% (4/22) to 36.4% (8/22) of mRNA-1273-vaccinated HCWs falling below detection limit for the four variants compared to 53.8% (14/26) to 73.1% (19/26) for BNT162b2 (**Fig. 2E**).

We examined additional factors that may contribute to strength and duration of the nAb response to vaccination, including age and sex. We observed no significant correlation for age and NT_50_ against D614G at any time point (**Fig. 3A-C**), potentially influenced by our relatively younger pool of study subjects. However, male HCWs exhibited significantly higher NT_50_ titers compared to females (**Fig. 3D**).

**Figure 3:**
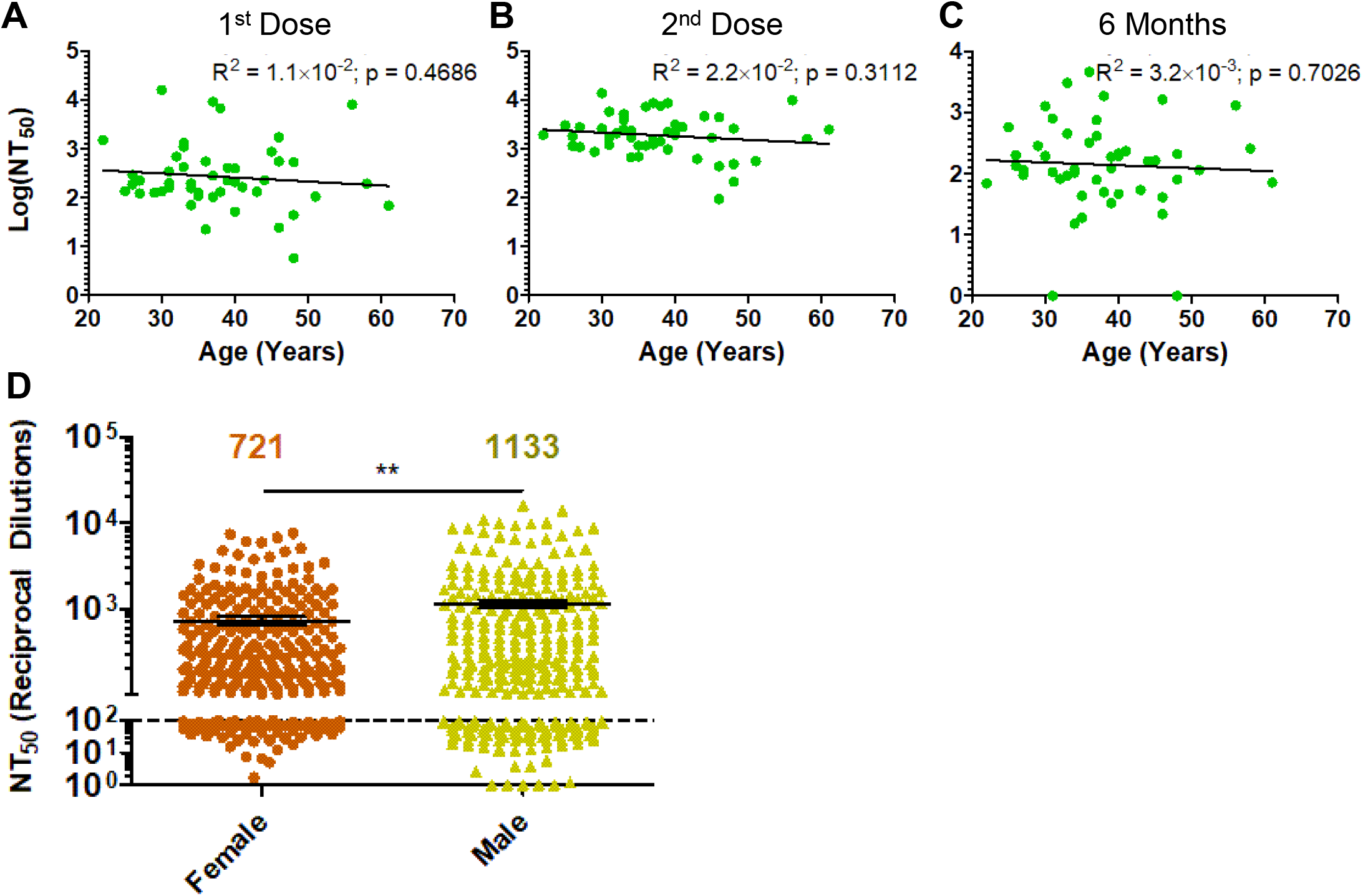
Impact of age and sex on response to mRNA vaccination. (**A-C**) Log_10_ transformed NT_50_ titers against D614G-SARS-CoV-2 S pseudotyped lentivirus are plotted against age (in years) at time of second vaccine dose for HCW samples collected post-fist vaccine dose (**A**), post-second vaccine dose (**B**), and 6 months post-second vaccine dose (**C**). (**D**) NT_50_ values against all variants at all time points were compared for male and female HCWs, with mean NT_50_ values displayed at the top of the plot. Statistical significance was determined by linear regression with least squares residual fit (**A-C**) or by unpaired, two-tailed student’s t-test with Welch’s correction (**D**). P-values are indicated as ‘ns’ (not significant) for p > 0.05 or *p < 0.05, **p < 0.01, ***p < 0.001.

## Discussion

In summary, we report a dramatic decline of SARS-CoV-2 nAb at six months post-mRNA vaccination and examined several key factors accounting for these kinetics. Most critically, we observed a drastic drop in nAb titers from 3-4 weeks to six months post-second vaccine dose, with more than 50% of HCWs exhibiting NT_50_ values below detection limit against Delta at the latter time. This number increased to almost 70% for the anti-N negative HCWs, which was in sharp contrast to that of anti-N positive HCWs, with 25% below the background. Thus, additional antigen exposures are necessary to improve the durability of the SARS-CoV-2 nAb response, consistent with data from administration of mRNA vaccine booster doses (16). Together, these results support a rationale for the need for boosters and alternative vaccination strategies to achieve long-term protection from infection with SARS-CoV-2.

Additionally, we observed that individuals vaccinated with BNT162b2 exhibited lower nAb titers than individuals vaccinated with mRNA-1273. However, the trend for declining nAb titers was consistent for both vaccines. Thus, both mRNA vaccines require booster doses to maintain protective nAb levels, although the waning of nAb responses likely occurs over a relatively longer period of time for mRNA-1273. Further examination of the durability of cellular immunity following mRNA vaccination is needed, as this more persistent immunity may limit the rates of hospitalization and death, which remain low for mRNA vaccinated individuals (17).

In this study, we found that all three VOCs consistently had reduced NT_50_ values compared to D614G at all time points, with Beta showing the most pronounced nAb resistance, followed by Delta. These results are consistent with preliminary reports from ours and other groups (14, 18, 19). However, we found that the Delta variant exhibited comparable or even higher resistance to nAbs than Beta for samples collected at six months post vaccination. The more modest drop in NT_50_ values at six months for the Beta variant was unclear, but likely the result of this variant’s pre-existing strong resistance to neutralization following the second dose of vaccination. Further, the more dramatic decline in nAb titers against Delta could be attributed to a lower frequency and durability of neutralizing antibody-producing plasma cells. As reported by others, the rampant spread of Delta in vaccinated and unvaccinated populations is likely related to other factors such as its high replication kinetics and transmissibility (20) coupled with its comparable neutralization resistance.

## Materials and Methods

### Health Care Worker Cohort

De-identified vaccinated health care worker (HCW)’s serum samples were collected under approved IRB protocols (2020H0228 and 2020H0527). These 48 HCWs ranged in age from 22-61 years (median = 37; IQR = 31.75-43.25) and included 26 male and 22 female HCWs. HCWs were vaccinated with either Moderna mRNA-1273 (n = 22) or Pfizer/BioNTech BNT162b2 (n = 26). Sera were collected from HCWs at 4 time points, with median time points being 222 days (IQR = 215-225.75) pre-first vaccine dose (Pre), 21 days (IQR = 19.25-23) post-first vaccine dose (Post 1^st^), 26 days (IQR = 22.5-28) post-second vaccine dose (Post 2^nd^), and 194 days (IQR = 190=197.75) post-second vaccine dose (6 Months). HCWs received their second vaccine dose between January and February of 2021.

HCW COVID-19 status was determined by anti-N ELISA (described below). Of the 48 HCWs examined, one was anti-SARS-CoV-2 N positive by ELISA pre-vaccination, four became anti-N positive for their post-first vaccine dose sample, three for their post-second vaccine dose sample, and four for their six-month vaccine sample—indicating that these 12 subjects were infected by SARS-CoV-2 at the different phases of vaccination.

### Constructs for Pseudotyping Virus Production

Production of lentiviral pseudotyped virus was performed using a previously reported protocol using pNL4-3-HIV-1-inGluc vector (13, 14, 21–23). This vector is a pNL4-3-HIV-1 ΔEnv construct and contains a *Gaussia* luciferase reporter gene with a CMV promoter both oriented in an anti-sense orientation relative to the HIV-1 genome. This *Gaussia* luciferase reporter gene then contains a sense orientation intron, which prevents expression of *Gaussia* luciferase in the virus producing cells. However, after the intron is spliced from full length virus genomes and upon integration into target cells, target cells can produce *Gaussia* luciferase, which is secreted in mammalian cell culture (24). Constructs encoding N- and C-terminal flag-tagged SARS-CoV-2 spike (S) for each variant — D614G, Alpha (B.1.1.7), Beta (B.1.351), and Delta (B.1.617.2) — were synthesized and cloned into pcDNA3.1 vector using KpnI/BamHI restriction enzyme cloning by GenScript BioTech (Piscataway, NJ).

### Cell Lines and Maintenance

HEK293T cells (CRL-11268, CVCL_1926, ATCC, Manassas, VA) and HEK293T-ACE2 cells (NR-52511, BEI Resources, ATCC, Manassas, VA) were maintained in Dulbeco’s Modified Eagles Medium (Gibco, 11965-092, ThermoFisher Scientific, Waltham, MA) supplemented with 10% (v/v) fetal bovine serum (F1051, Sigma-Aldrich, St. Louis, MO) and 1% (v/v) penicillin/streptomycin (SV30010, HyClone Laboratories Inc., Logan, UT). Cells were maintained in at 37°C and 5% CO_2_.

### Pseudotyped Virus Production and Titering

Pseudotyped lentivirus was produced by co-transfection of HEK293T cells with pNL4-3-HIV-1-inGluc and pcDNA3.1 vector expressing the spike of interest (D614G, B.1.1.7, B.1.351, or B.1.617.2) in a 2:1 ratio using polyethylenimine (PEI) transfection. Virus was collected 24 hrs, 48 hrs, and 72 hrs after transfection, then was pooled and stored at −80°C.

To determine relative titers of harvested virus, the pseudotyped virus for each of the SARS-CoV-2 variants were used to infect HEK293T-ACE2 cells. Then, 48 hrs and 72 hrs after infection, *Gaussia* luciferase activity in the media of infected cells was determined. 20 μL of cell culture media and 20 μL of *Gaussia* luciferase substrate (0.1M Tris (T6066, MilliporeSigma, Burlington, MA) pH 7.4, 0.3M sodium ascorbate (S1349, Spectrum Chemical Mfg. Corp., New Brunswick, NJ), 10 μM coelenterazine (CZ2.5, GoldBio, St. Louis, MO)) were combined in a white polystyrene 96-well plate. Luminescence was immediately measured by a BioTek Cytation5 plate-reader.

### Virus Neutralization Assays

Virus neutralization assays were performed as previously reported (13, 14, 23). In a 96-well format, HCW serum was 4-fold serial diluted and 100 μL of pseudotyped virus was added (final dilutions of 1:80, 1:320, 1:1280, 1:5120, 1:20480, and no serum). Note that, to ensure comparable results between SARS-CoV-2 variants, equivalent amounts of infectious virus were used based on the pre-determined virus titers. The virus was incubated with HCW serum for 1 hr at 37°C, followed by infection of HEK293T-ACE2 cells seeded on a 96-well polystyrene tissue culture plate. *Gaussia* luciferase activity in cell culture media was then assayed 48 hrs and 72 hrs after infection as described above. Neutralizing titer 50% (NT_50_) for each serum sample was determined by non-linear regression with least squares fit in GraphPad Prism 5 (GraphPad Software, San Diego, California).

### Anti-N ELISA

Anti-N ELISA was performed as previously reported (13). ELISA was performed by using the EDI Novel Coronavirus COVID-19 N protein IgG ELISA Kit (KT-1032, EDI, San Diego, CA) following manufacturer’s protocol. Briefly, 100 μL of a 1:100 dilution of HCW serum was added to microplates coated with SARS-CoV-2 neucleocapsid (N) antigen and incubated for 30 min. Plates were then washed and treated with 100 μL of HRP labeled anti-human-IgG antibody (31220, EDI, San Diego, CA) for 30 min. Then plates were washed and 100 μL of ELISA HRP substrate (10020, EDI, San Diego, CA) was added and incubated for 20 min before 100 μL of stop solution (10030, EDI, San Diego, CA) was added. Absorbance at 450 nm was read by spectrophotometric plate reader using Gen 5 software.

### Statistical Analyses

Statistical analysis was done with GraphPad Prism 5. Comparisons between multiple groups were done using one-way ANOVA with Bonferroni’s multiple testing correction (Figs. 1A, 2B) or one-way repeated measures ANOVA with Bonferroni’s multiple testing correction (Figs. 1C-F). For comparisons between two “treatments” across multiple groups, a two-way ANOVA with Bonferroni’s multiple testing correction was used (Figs. 2C and 2E). For comparisons between two groups, an unpaired, two-tailed student’s t-test with Welch’s correction was used (Figs. 2D, S1D). For correlative analyses between two continuous variables, a linear regression model with least squares fit was used with log_10_ transformed NT_50_ values to better approximate linearity (Figs. 1G-J, S1A-C).

## Author Contributions

J.P.E. conducted neutralization assays, analyzed data, and drafted the manuscript. C.Z. aided in neutralization assays, review of manuscript, and provided valuable discussion and insight. C.C. contributed to recruitment of HCWs and sample collection. R.J.G. contributed to study design, provided HCW samples and subject information, reviewed the manuscript, and provided valuable discussion and insight. G.L. provided anti-N ELISA data. S.-L.L. contributed to study design, directed laboratory personnel, and aided in drafting and revision of the manuscript. C.C., G.L., L.J.S., E.M.O., and R.J.G. provided valuable discussion and revision of the manuscript.

## Acknowledgements

We thank David Derse, Marc Johnson, Fang Li, and Ali Ellebedy for providing plasmids, cells, and antibodies. We thank members of Liu lab for sharing reagents and discussion. We also thank the NIH AIDS Reagent Program and BEI Resources for supplying important reagents that made this work possible. We thank the Clinical Research Center/Center for Clinical Research Management of The Ohio State University Wexner Medical Center and The Ohio State University College of Medicine in Columbus, Ohio, specifically Francesca Madiai, Claire Carlin, Dina McGowan, Breona Edwards, Evan Long, and Trina Wemlinger, for logistics, collection and processing of samples.

